# Genomic signatures distinguish isolates of *Salmonella* Typhimurium from humans and livestock

**DOI:** 10.64898/2026.11.05.764597

**Authors:** Giovanni Merici, Erika Scaltriti, Stefano Pongolini, Martina Tambassi

## Abstract

*Salmonella enterica* serovar Typhimurium is a major cause of human salmonellosis and livestock represent an important reservoir for human infection through the food chain. However, it remains unclear whether isolates recovered from human disease reflect the broader livestock population or constitute a subset of livestock-associated isolates characterized by specific genetic features.

Here, we analysed 1,398 *S.* Typhimurium genomes from human, poultry, bovine and swine isolates to evaluate whether isolates associated with human disease can be distinguished from those circulating in livestock and to identify the genetic features underlying this discrimination.

Whole-genome multilocus sequence typing profiles were converted into a gene presence/absence matrix and analysed using four machine-learning algorithms, which showed high and consistent discriminatory performance. Model performance was influenced by the phylogenetic structure of the population and by the variable distribution of human- and livestock- associated isolates across lineages of sequence type (ST) 19, the most prevalent S. Typhimurium ST. Nevertheless, within ST19, 97 genes were significantly enriched and 36 were significantly depleted in human-associated genomes relative to genomes from each of the three livestock sources.

Human-enriched genes included 14 genes required for biosynthesis of the lipopolysaccharide core, the O-antigen, the T1 antigen, and six virulence factors. These genes were consistently absent from livestock-associated genomes across ST19 lineages. Conversely, human-depleted genes included the galactitol utilisation cluster and five virulence genes, which are predominantly absent in specific ST19 lineages. Overall, our findings support the existence of host-associated genomic signatures that distinguish isolates associated with human disease from those sampled from the livestock reservoir.

**Data Summary:** This study analysed publicly available *Salmonella* Typhimurium genomes; no new sequencing data were generated. Genome assemblies, associated metadata and whole-genome multilocus sequence typing (wgMLST) allelic profiles were retrieved from EnteroBase (https://enterobase.warwick.ac.uk). The final dataset comprised 1,398 genomes, as detailed in Table S1. Custom analysis scripts, input datasets, intermediate files and analysis outputs are available in the GitHub repository (https://github.com/GiovanniMerici/Typhimurium_Human_vs_Animal). These include the train/test split membership, selected model hyperparameters and per-split SHAP matrices. Software versions and analysis parameters are described in the Methods.

**Impact statement:** *Salmonella enterica* serovar Typhimurium is among the leading causes of human salmonellosis, with livestock representing important reservoirs of infection. Genomic studies have primarily focused on identifying the animal reservoirs associated with human clinical isolates, while whether these isolates represent the broader bacterial populations circulating in livestock or a genetically distinguishable subset remains comparatively underexplored. Using 1,398 genomes and four machine-learning models, we demonstrate that human-associated isolates can be distinguished from poultry, bovine, and pig isolates with high accuracy and reproducibility. Although phylogenetic structure contributed to the classification, we were able to identify specific genes significantly enriched and depleted in human-associated genomes relative to genomes from each of the three livestock sources within the most prevalent S. Typhimurium ST. Importantly, these genomic differences were biologically coherent, rather than scattered across unrelated loci, involving genes associated with surface-antigen biosynthesis and virulence. These findings extend genomic source-attribution approaches by showing that human clinical isolates represent a distinguishable subset of livestock-associated *S*. Typhimurium populations. This distinction provides a complementary perspective for interpreting genomic surveillance data and highlights the potential relevance of host-associated genomic variation for understanding the relationship between livestock reservoirs and human infection.

## Introduction

Non-typhoidal *Salmonella* (NTS) is a major cause of foodborne disease worldwide, resulting in substantial morbidity and representing a persistent public health burden.^1^ NTS circulates among humans, animals, and the environment, with livestock representing major reservoirs for human infection through the consumption of animal-derived foods. However, whether NTS strains responsible for human disease are broadly representative of the *Salmonella* population circulating in livestock and entering the food chain, or whether they represent a non-random subset enriched for specific genomic features that may facilitate their ability to cause human disease remains to be determined. This question is particularly relevant given that host- associated diversification has been documented in *Salmonella*, with certain serovars and intraserovar lineages exhibiting genomic changes associated with adaptation to specific animal hosts.^2,3^

Among NTS, *Salmonella enterica* serovar Typhimurium is one of the main serovars associated with human disease. According to surveillance data from the U.S. Centers for Disease Control and Prevention (CDC) and, in the European Union, from the European Food Safety Authority (EFSA) and the European Centre for Disease Prevention and Control (ECDC), *S.* Typhimurium is the third most frequently reported *Salmonella* serovar causing human disease in the United States and the second most frequently reported serovar in the European Union.^4,5^

*S.* Typhimurium can infect humans as well as a wide variety of animal hosts, including all the main livestock species.^4^ Within the most common *S*. Typhimurium sequence type (ST19), several lineages associated with specific animal hosts have been identified, providing a clear example of the host-associated diversification observed in *Salmonella*.^6^

The epidemiological importance and broad host range of *S.* Typhimurium make it an ideal model to investigate whether strains associated with human disease carry genomic features that distinguish them from strains circulating in livestock. Because this serovar infects multiple livestock species, such comparisons can be performed across a diverse animal population, reducing the likelihood that the identified genomic features simply reflect adaptation to a particular animal host.

Whole-genome sequencing and machine-learning approaches provide powerful tools for identifying genetic features that characterize a specific group of strains. While these approaches have been successfully employed in *Salmonella* genomics to address diverse epidemiological and evolutionary questions, such as the discrimination of typhoidal and non-typhoidal strains and source attribution analyses, their potential for identifying genomic signatures associated with strains causing human disease remains largely unexplored.^7–10^

In this study, we applied machine-learning approaches to 1,398 genomes of *S.* Typhimurium isolates from human, poultry, bovine and swine sources collected in North America between 2020 and 2025, to investigate the genetic features associated with human-associated isolates. We used whole-genome allelic profiles to assess variation in gene presence and predicted gene integrity, accounting for both gene loss and potential gene inactivation, to identify genomic features associated with human-derived isolates. By integrating machine-learning feature importance with phylogenetic information, we discriminated lineage-specific signals from genetic features consistently associated with human-associated isolates, thereby highlighting candidate determinants with broader relevance to *Salmonella* epidemiology. Our findings provide new insights into the genomic features associated with human-associated *S.* Typhimurium and offer a framework for identifying genetic factors that may contribute to the successful transmission of strains associated with human disease.

## Methods

### Dataset Construction and Balancing

The genomes used in this work were retrieved from EnteroBase (*Salmonella* database; https://enterobase.warwick.ac.uk).^11^ Specifically, we selected genomes of *S.* Typhimurium strains isolated in North America between 2020 and 2025 and classified as human, poultry, bovine, or swine sources. The poultry, bovine, and swine categories included isolates obtained directly from animals as well as from food products derived from the corresponding animal source. Strain metadata, genome assembly, assembly information, wgMLST scheme identifiers and wgMLST allelic profiles were collected for 5,954 *Salmonella* Typhimurium strains originating from poultry (2,607), cattle (696), swine (1,552), and humans (1,099). The dataset was then manually curated as follows: in case of duplicated wgMLST identifiers a single representative record was retained; genomes with source details not matching the source type were removed. In silico serotyping was performed on genome assemblies using Salmonella In Silico Typing Resource (SISTR, version 1.1.3) with quality control enabled (sistr --qc -f csv).^12^ Only S. Typhimurium genomes satisfying the following criteria were retained: SISTR serovar prediction equal to "Typhimurium", phase 1 flagellar equal to antigen i, phase 2 flagellar equal to antigen 1,2, and SISTR QC status PASS. This step excluded monophasic and antigenically atypical variants, retaining 2,340 poultry, 411 bovine, 383 swine and 699 human genomes. Finally, the dataset (3833 genomes) was balanced across human vs animal sources by random sampling without replacement using a fixed seed (random_state = 42, pandas 2.2.2). Specifically, all 699 confirmed human genomes were retained while 233 isolates were drawn from each animal source (poultry, bovine and swine), giving a final dataset of 1,398 genomes (Table S1).

### SNP-based phylogeny

Core Single-Nucleotide Polymorphisms (core-SNPs) were identified from genome assemblies through Snippy v4.6.0 (https://github.com/tseemann/snippy) in contig mode (snippy --ctgs) using *S*. Typhimurium LT2 genome (GCF_000006945.2, ASM694v2) as reference. A core-genome alignment was generated with snippy-core, cleaned with snippy-clean_full_aln and recombinant regions were predicted and masked with Gubbins v3.1.0 (run_gubbins.py, 16 threads), which converged after one iteration.^13^ Constant-site counts were obtained from the cleaned alignment with snp-sites -C and supplied to IQ-TREE v3.0.1 for ascertainment-bias correction (-fconst).^14,15^ A maximum-likelihood phylogeny was inferred from the recombination-filtered polymorphic-site alignment with the best-fit substitution model selected by ModelFinder under BIC (-m MFP), with branch support assessed by 1,000 ultrafast bootstrap replicates (-bb 1000) and 1,000 SH-aLRT replicates (-alrt 1000).^16^ The phylogenetic tree was rooted using ST36 genomes as the outgroup due to their genetic divergence from other S. Typhimurium STs.^17^ The tree was visualised and annotated in iTOL v6.^18^

### Generation of the functional gene presence/absence matrix

The wgMLST allelic profiles downloaded from EnteroBase were converted into a functional gene presence/absence matrix.^11^ To this end, the allele sequences of the wgMLST loci, each representing a gene, were screened for the presence of internal stop codons with Biopython 1.87 and flagged as truncated if TAA, TAG or TGA occurred before the final codon.^19^ For each truncated allele, the position of the first internal stop codon and the implied truncated protein length were retained, and the allele number was recovered from the allele identifier, producing an explicit list of locus–allele combinations predicted not to encode a full-length product. Truncated alleles were considered non-functional and coded as 0, along with missing and invalid alleles. All other alleles were considered functional and coded as 1. Non-informative loci (i.e., loci showing no variation in functional status across all genomes) were removed. The resulting binary profiles were used to construct a functional gene presence/absence matrix.

### Machine learning analyses

#### Model comparison

The functional gene presence/absence matrix was subjected to supervised machine learning approaches. To this end, the binary profiles were assigned to either the ‘human’ or ‘animal’ category according to the isolation source of the corresponding strains. Four algorithms were used and compared: L2-penalised logistic regression (saga solver, max_iter = 5000, class_weight = "balanced", preceded by StandardScaler (with_mean = False) in a scikit-learn pipeline), random forest (500 trees, min_samples_leaf = 2, class_weight = "balanced"), extremely randomised trees (also called Extra Trees, same settings applied to random forest), and gradient-boosted trees (XGBoost 3.3.0; 500 estimators, max_depth = 4, learning_rate = 0.03, subsample = 0.8, colsample_bytree = 0.8, binary:logistic objective, hist tree method).^20,21^ All models used random_state = 42.

Models were evaluated over 50 independent iterations, with the dataset randomly split into stratified training (80%) and test (20%) sets at each iteration. The stratification was based on the binary human/animal label to preserve class proportions in each partition. Therefore, each split comprised 1,118 training and 280 test genomes (559 and 140 per class, respectively). The same data partitions were used for all four algorithms to ensure that differences in performance were attributable to the algorithms themselves rather than to differences in the data partitions.

Performance was summarised as the mean and standard deviation across splits of ROC-AUC, accuracy, balanced accuracy, F1 score, precision, and recall.

#### Sensitivity analysis

To assess whether the obtained discrimination between human and animal-derived genomes was influenced by STs, XGBoost was applied using as features (i) ST, (ii) the functional gene presence/absence profiles of ST19 genomes, (iii) all the functional gene presence/absence profiles except those belonging to ST36 genomes, and (iv) five random subsets of the full dataset matching to the size and source balance of the ST19 cohort. For these analyses, the same 50 stratified 80/20 train/test splits and the same fixed XGBoost hyperparameters described above were used. Loci that were invariant within a given cohort were removed before model fitting.

To assess whether the obtained discrimination between human and animal-derived genomes was influenced by ST19 clusters, patristic distances between the ST19 genomes were computed from the recombination-filtered maximum-likelihood phylogeny and used for single- linkage hierarchical clustering, with clusters defined at seven thresholds from 1 × 10⁻ to 1 × 10⁻ substitutions per site, yielding from 1,184 to 15 clusters. Models were evaluated by stratified group cross-validation (five folds, ten repeats, giving 50 evaluations as in the above analyses), in which all genomes of a cluster were assigned to either the training or the test partition. Folds in which either partition contained only human-derived or animal-derived genomes were skipped.

### XGBoost Model Tuning and SHAP-Based Feature Attribution

XGBoost was re-evaluated using a nested resampling framework to combine hyperparameter tuning with unbiased performance estimation and feature attribution (**Fig S1**). The outer loop consisted of the same 50 stratified 80/20 splits described above. Each split comprised 1,118 training genomes (559 human and 559 animal) and 280 held-out test genomes (140 human and 140 animal). Within the training partition of each outer split, hyperparameters were tuned by randomised search (RandomizedSearchCV, 20 sampled configurations, 3-fold stratified inner cross-validation, ROC-AUC as the selection criterion, seed 42 + split_id), giving 60 model fits per outer split. The search space covered n_estimators (200–700), max_depth (2–6), learning_rate (0.01–0.1), subsample (0.6–1.0), colsample_bytree (0.5–1.0), min_child_weight (1–10), gamma (0–1.0), reg_alpha (0–1.0) and reg_lambda (0.5–5.0). The best configuration was refitted on the full training partition and evaluated on the held-out isolates of that outer split. Best hyperparameters, per-split performance and split membership were recorded for every outer split.

A SHAP-based feature-importance analysis was then performed to identify the genes contributing most to the tuned XGBoost predictions.^22^ SHAP values were computed with shap.TreeExplainer 0.52.0 on the held-out binary profiles of each outer split. For each split and gene, the mean, median and maximum absolute SHAP value and the signed mean SHAP value were recorded, and genes were ranked within the split by mean absolute SHAP value. Across the 50 splits, per-gene statistics were aggregated into a mean, standard deviation and median of the per-split mean absolute SHAP value, together with the mean, median and best within-split rank. Rank stability was quantified as the number and proportion of splits in which a gene appeared among the top 10, 25, 50 and 100 loci. Genes were finally ordered by mean absolute SHAP value, breaking ties on top-50 frequency and mean rank. Genes with a non-zero SHAP contribution in at least one split (defined as SHAP-ranked genes, n = 1,774) were carried forward to functional annotation.

### Functional annotation of SHAP-ranked genes

Three complementary annotation strategies were then applied to the SHAP-ranked genes. The functional description associated with each wgMLST locus in the EnteroBase *Salmonella* scheme was retrieved together with the corresponding locus tag and, where available, the gene symbol.^11^ Then, orthology-based functional annotation was performed with eggNOG-mapper v2.1.15 (emapper.py --itype CDS -m diamond --tax_scope Bacteria --cpu 8) on the reference allele (allele 1 of the wgMLST) of each locus retrieved from EnteroBase.^11^ DIAMOND was used for the seed-orthologue search against the eggNOG v5.0 database and orthologue assignment was restricted to the Bacteria taxonomic scope. ^23,24^ For each locus, the seed orthologue, eggNOG orthologous groups, COG functional category, free-text description, preferred gene name, and GO, EC, KEGG and PFAM identifiers were retained.

Finally, the nucleotide sequence of each gene was translated using the bacterial genetic code (NCBI translation table 11) in Biopython, with alternative bacterial start codons ATG, GTG, TTG resolved to methionine. Sequences that failed strict CDS validation were translated permissively and flagged accordingly, and translation status was retained for every gene.^19^ Translated sequences underwent structure-informed functional annotation with Baktfold v0.1.0 (baktfold proteins -t 4 --foldseek-gpu). Baktfold uses the ProstT5 protein language model to convert amino-acid sequences into the foldseek 3Di structural alphabet and then searches against Swiss-Prot, the AlphaFold Database cluster representatives, the Protein Data Bank and CATH.^25,26^

A locus was considered annotated when EnteroBase or Baktfold returned a product other than "hypothetical protein", and when eggNOG-mapper assigned an orthologous group with a COG functional category other than S (function unknown).

### Identification of human-enriched and human-depleted SHAP- ranked genes

For each ST, genes were classified as human-enriched when their presence frequency was significantly higher in genomes of human origins than in genomes from animal origins. Conversely, genes were classified as human-depleted when their presence frequency was significantly lower in genomes from humans than in genomes from animal origins. Specifically, for ST36, ST213, ST34, and ST2072, the presence frequency of SHAP-ranked genes in genomes of human origin was compared to that in genomes of animal origin using Fisher’s exact test. For ST19, the presence frequency of SHAP-ranked genes in genomes of human origin was compared separately with the corresponding frequencies in genomes of poultry, bovine, and swine origin, and the obtained *p*-values from the Fisher’s exact tests were adjusted for multiple comparisons using the Bonferroni correction.

### Genomic localisation and spatial clustering

To test whether human-enriched and human-depleted genes were physically clustered in the genome, their reference sequences were mapped to the *S.* Typhimurium LT2 genome (chromosome NC_003197.2 and plasmid NC_003277.2; assembly GCF_000006945.2, ASM694v2). A nucleotide BLAST database was built with makeblastdb and searches were run with BLASTN 2.12.0 (-evalue 1e-5, -max_target_seqs 5, tabular output including query and subject coordinates, alignment length, percentage identity, bit score, e-value and query length).^27^ Hits were retained only if they showed at least 90 % nucleotide identity and at least 80 % query coverage, and a single best hit was kept per gene by ordering on coverage, percentage identity, bit score and e-value. Genomic start and end coordinates were defined as the minimum and maximum of the subject coordinates, and strand was inferred from their order.

Distances between neighbouring genes were computed within each replicon after sorting by genomic coordinate, as the start coordinate of the next gene minus the end coordinate of the current gene, with overlapping genes assigned a distance of zero. Loci on the same replicon separated by no more than 10 kb were assigned to the same spatial cluster.

Only groups of two or more loci were considered clusters. Spatial clustering was assessed against a null model in which the same number of genes was placed uniformly at random along the LT2 chromosome (4,857,450 bp) and grouped under the same 10 kb criterion, repeated 20,000 times with a fixed seed. P values are the proportion of permutations in which the number of loci falling within clusters, or the size of the largest cluster, reached or exceeded the observed value.

## Results

### *S*. Typhimurium population structure

The dataset generated in this study included genomes of 1,398 *S*. Typhimurium isolated in North America, from humans, poultry, bovine, and swine (**Fig. 1**). The dataset was balanced to include an equal number of genomes of isolates from human and livestock sources. Most isolates originated from the United States (70.5%), followed by Canada (27.3%) and Mexico (2.2%). The five most common sequence types were ST19 (89.2%), ST36 (5.36%), ST213 (1.72%), ST34 (1.22%), and ST2072 (1%). Source distributions varied across STs. ST19 comprised similar proportions of human- and livestock associated genomes (47.2% human, 16.1% swine, 18.1% bovine, and 18.5% poultry genomes). ST36 was predominantly of human origin, with 98.7% of genomes derived from humans and 1.3% from swine. ST34 comprised mainly swine-associated genomes (58.8%) with the remaining 41.2% of human origin. ST213 comprised 8.33% human-, 79.1% swine-, and 12.5% bovine-associated genomes, whereas ST2072 comprised 64.29% human-, 7.14% poultry-, 14.29% swine-, and 14.29% bovine- associated genomes.

**Figure 1.**
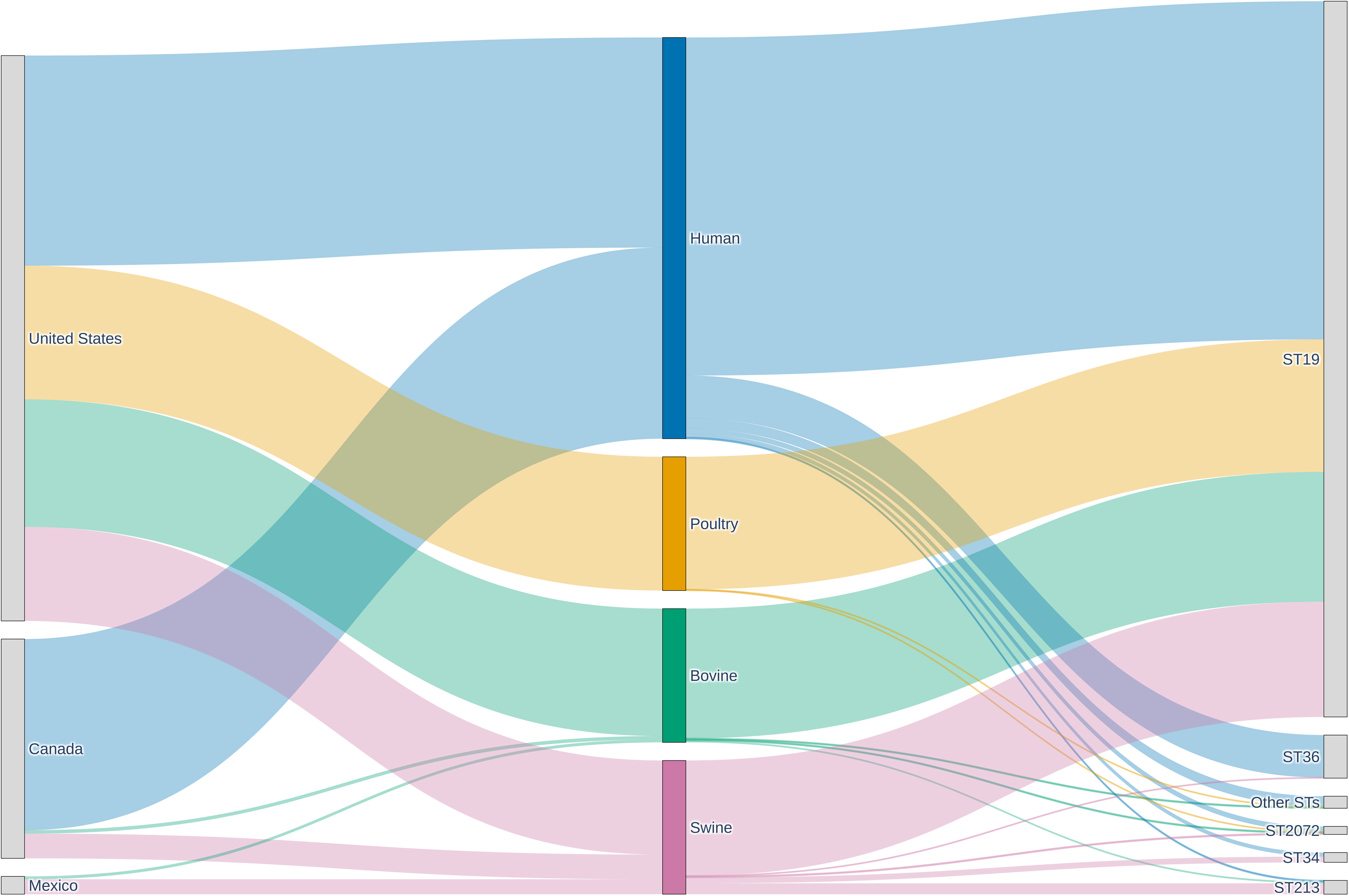
Dataset composition. Sankey diagram representing the country of isolation, the source type and the five most common STs of the *S*. Typhimurium isolates included in our dataset.

To characterize the dataset structure and assess the relatedness of human and livestock- associated genomes, we performed a core SNP-based phylogenetic analysis (**Fig. 2**). The phylogenetic tree was rooted on genomes belonging to ST36, because genetically distinct to the other main *S*. Typhimurium STs.^17^ As expected, the phylogenetic analysis revealed a distinct lineage comprising the ST36 genomes. In contrast, ST34, ST213, and ST2072 formed distinct lineages within ST19, highlighting their phylogenetic diversification within this lineage. Focusing on genomes belonging to ST19, these were divided into distinct sublineages, some of which were predominantly composed of human-derived genomes, whereas others were mainly associated with poultry, bovine and swine sources. This pattern is consistent with previous studies reporting host-associated population structure within S. Typhimurium ST19.^6^ However, human-associated genomes were generally intermingled with livestock-associated genomes, as previously observed.^8^ Overall, these results support the representativeness of our dataset with respect to the known population structure of *S*. Typhimurium and indicate that human- and livestock-associated isolates cannot be readily distinguished based on their core SNP-based phylogenetic relationships.^9,17^

**Figure 2.**
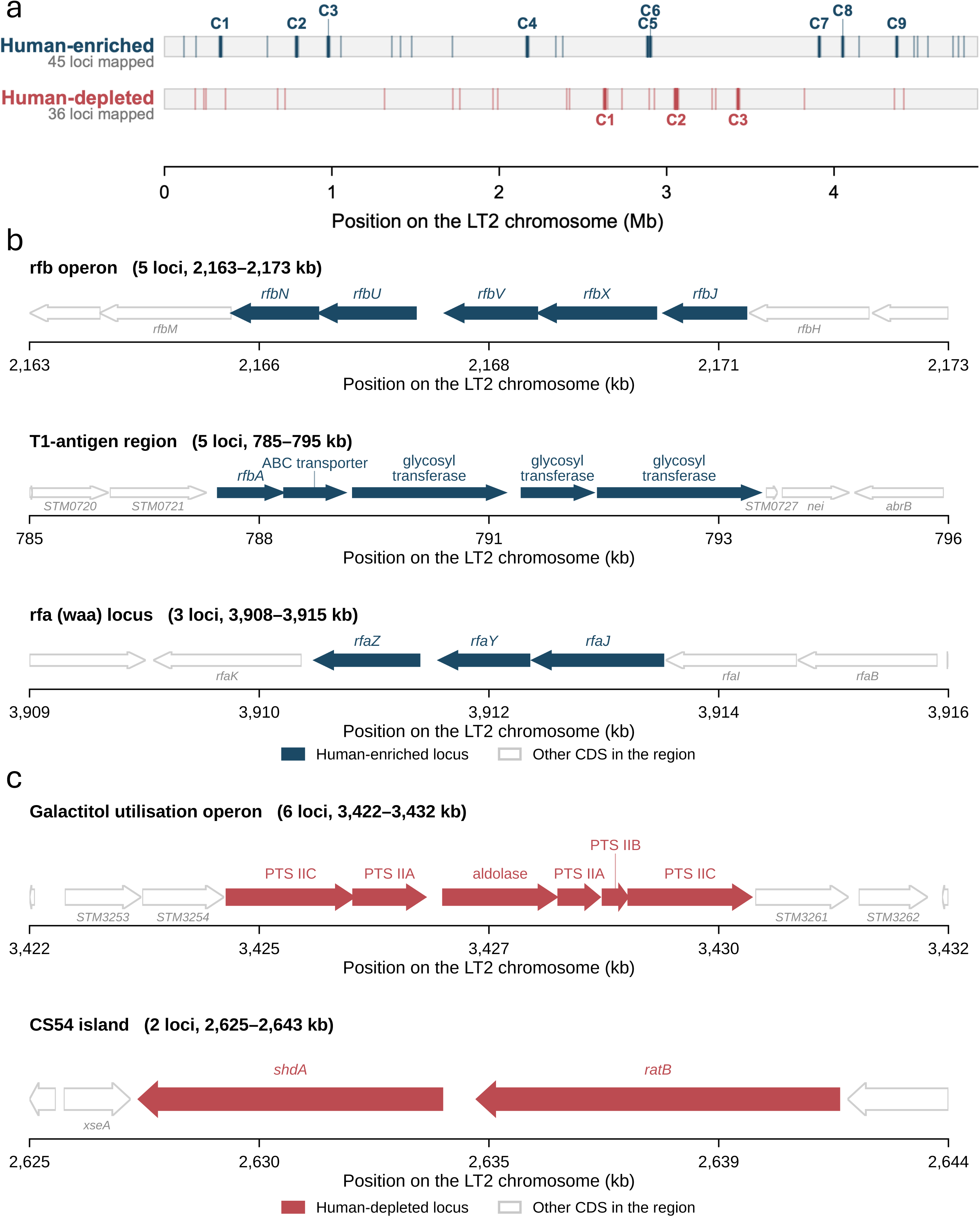
Core-SNP phylogenetic analysis of the studied *S*. Typhimurium genomes. Innermost annotation ring specifies source type; second annotation ring specifies the five most common STs.

### Applying machine learning to discriminate between human- and livestock-associated genomes

As core SNPs phylogeny did not clearly discriminate between human- and livestock-associated genomes, we investigated whether the analysis of gene gain/loss or pseudogenization events could provide discriminatory information. To this end, the wgMLST allelic profiles, encompassing both core and accessory genes, were converted into a functional gene presence/absence matrix by coding alleles that were absent, truncated, or contained premature stop codons as “absent,” and all other alleles as “present.” The obtained matrix, including a total of 6,254 genes, was used as input in machine learning approaches. Four supervised machine- learning algorithms, L2-penalised logistic regression, Random Forest, ExtraTrees, and XGBoost, were applied and compared using the same 50 repeated stratified train/test splits, preserving the human/animal proportions in each split. All four models showed high and relatively stable discriminatory performances across the 50 repeated data splits, indicating that genomes of human and animal origin differ according to their functional gene profiles (**Fig. 3a**). XGBoost achieved the highest mean ROC-AUC (0.949 ± 0.010), balanced accuracy (0.885 ± 0.018), and F1 score (0.886 ± 0.018) and was therefore selected for subsequent analyses.

**Figure 3.**
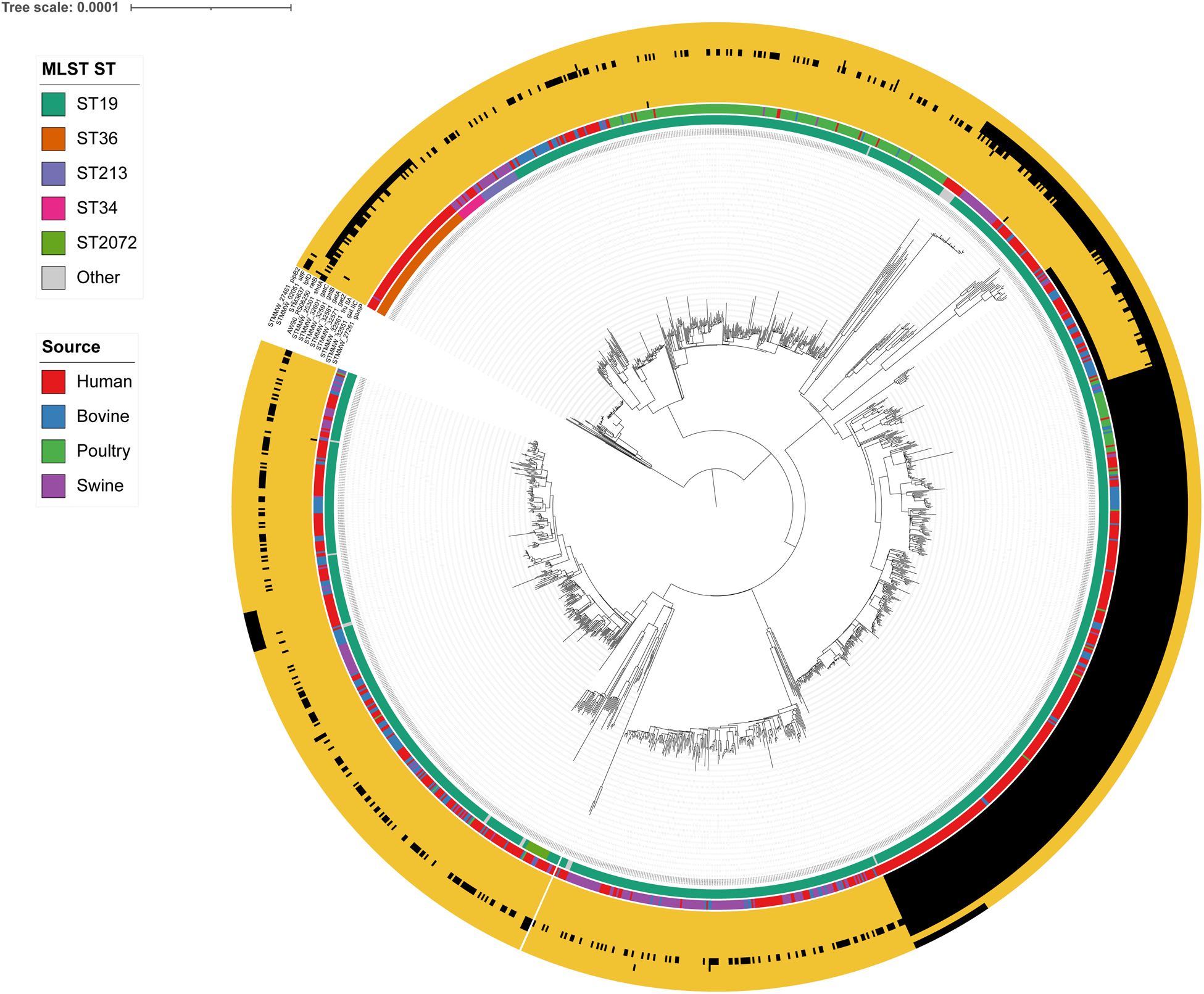
Model performance and sensitivity analyses. **(a)** Performance of four models for human-versus-animal source attribution using wgMLST presence–absence data. Bars show means across 50 stratified train–test splits; error bars represent SD. **(b)** Sensitivity of human- versus-animal discrimination to sequence type. ROC-AUC across 50 stratified train/test splits; boxes span the interquartile range, points are individual splits, means are given above. Loci invariant within a cohort were removed before fitting, so feature counts differ. The dashed line marks random expectation. **(c)** Discrimination within ST19 under cluster-aware evaluation. Whole clusters of related genomes, defined by patristic distance on the phylogeny, were assigned entirely to the training or the test partition. Boxes span the interquartile range, points are individual evaluations, means are given above; folds in which either partition contained a single class were skipped, so the number of evaluations decreases at coarser thresholds.

We next assessed whether the ability of machine-learning models to distinguish between human- and livestock-associated genomes reflected genetic features genuinely associated with the source of isolation or was instead influenced by phylogenetic differences among STs and by their diverse distribution across the two sources (**Fig. 3b**). A classifier trained on STs instead of functional gene presence/absence matrix showed limited predictive performance (ROC-AUC 0.577 ± 0.016), suggesting that ST identity alone was insufficient to explain the discrimination between the two sources. Differently, performance was unchanged when the analysis was restricted to ST19, which includes similar proportions of human- and livestock-associated genomes (0.951 ± 0.011), and when ST36— phylogenetically distant from the other STs and comprising almost exclusively human isolates—was excluded (0.945 ± 0.011). Furthermore, a random subset of the full dataset matching the size and human/animal proportions of ST19 gave 0.947 ± 0.011, indicating that the dataset size does not affect performance. The observed discrimination between genomes of human and animal origin was therefore not fully explained by ST-specific genetic features and distribution across sources.

The phylogeny also revealed substantial substructure within the most represented ST19, with genomes clustering into distinct lineages that can be predominantly associated with a specific source. Therefore, we further assessed whether the obtained discrimination between human- and livestock-associated genomes was influenced by the genetic differences between ST19 lineages. To this end, ST19 genomes were clustered using seven thresholds (1 × 10⁻ – 1 × 10⁻ substitutions per site) applied to their pairwise patristic distances. These distances quantify the evolutionary divergence between genomes along the phylogenetic tree, with increasing thresholds grouping progressively more distantly related genomes into fewer, larger clusters. Each cluster was assigned exclusively to either the training or test partition to assess whether the discrimination between human- and livestock-associated genomes generalized across ST19 clusters, rather than relying on cluster-specific genetic features. When closely related genomes were allowed to occur in both partitions (i.e. when a lower threshold was applied), performance remained high, whereas restricting entire, increasingly divergent lineages to either the training or test partition (i.e. when a higher threshold was applied) reduced predictive performance. This effect was reflected in a progressive decrease in ROC-AUC, from 0.949 ± 0.011 with 1,184 clusters, to 0.657 ± 0.135 with 15 clusters (**Fig. 3c**). These results indicate that the ability of the model to discriminate between human- and livestock-associated genomes was partly dependent on phylogenetic relatedness between training and test clusters. This suggests that part of the predictive signal was derived from lineage-specific genetic features, potentially reflecting the predominance of certain lineages in specific sources.

### Functional analysis of genes enriched and depleted in human- derived genomes

We next analysed which genes contributed most to the discrimination between human- and livestock-associated genomes. To this end, XGBoost was first re-evaluated using a nested resampling framework based on the same 50 stratified splits described above, to provide robust estimates of model performance and feature importance. The tuned XGBoost model achieved a mean held-out ROC-AUC of 0.948 ± 0.010, and a mean balanced accuracy of 0.882 ± 0.018 (**Fig S2**). These values were comparable to those obtained during the initial model comparison, indicating that model performance was stable across different train/test partitions and was not dependent on a single favorable split.

A SHAP-based feature-importance analysis was then performed to identify genes contributing most to the XGBoost predictions. Of the 6,254 genes included in the gene presence/absence matrix, 1,774 showed a non-zero aggregated SHAP importance, indicating that only a subset of genes contributed detectably to the model predictions (Table S2). The SHAP-ranked genes were annotated using the curated functional descriptions available in EnteroBase, the orthology- based annotation obtained with eggNOG-mapper, and the structure-informed annotation obtained with Baktfold.^11,23,26^ Considering the three approaches together, 1,546 loci (87.1%) were annotated by at least one source (**Fig. S3**, Table S3).

Finally, for each of the five most common STs, we identified genes occurring in a significantly higher proportion in human- compared to livestock-associated genomes, defined as human- enriched genes, and loci showing the opposite pattern, defined as human-depleted genes (Table S4). ST36 was excluded from the analysis because only one genome of animal origin was present in the dataset, precluding a robust comparison of human- and livestock-associated genomes belonging to this ST. A total of four genes encoding RfbV and RfbX, involved in biosynthesis of lipopolysaccharide (LPS) O-antigen, the virulence factor SpvD, and a putative ATPase resulted enriched in ST2072 human genomes, whereas no human-depleted genes were found. Regarding ST34, the analysis identified three human-enriched and six human- depleted genes. The human-enriched genes include those encoding the sulfonamide resistant dihydropteroate synthase Sul2 and the β-lactamase TEM-1, whereas among the human- depleted genes we found that encoding the host colonisation factor ShdA. No human-enriched or human-depleted genes were found for ST213.

Regarding ST19, a host-stratified analysis was performed to distinguish signatures associated with human origin from those specific to lineages associated with a particular host. Gene frequencies were therefore calculated separately for human-, poultry-, bovine-, and swine- derived genomes. Genes were considered human-enriched or human-depleted only if they occurred at a significantly higher or lower proportion, respectively, in human-associated genomes than in genomes associated with each of the three animal groups. A total of 97 genes were found to be human-enriched, whereas 36 were human-depleted. The two gene sets were analysed for their function and their genomic localization. Human-enriched set includes 14 genes involved in the biosynthesis of LPS, a major outer-membrane endotoxin and virulence factor of Gram-negative bacteria. Specifically, genes required for the LPS core (*rfaZ*, *rfaY*, *rfaJ*), O-antigen (*rfc*, *rfbN*, *rfbU*, *rfbV*, *rfbX*, and *rfbJ*), and T1 antigen (STM0722- STM0726) biosynthesis were found.^28,29^ The frequency of these genes was, on average, 18% ± 10% higher in human- than in livestock-associated genomes, with some livestock-associated genomes lacking only a subset of these genes, while others lacked them entirely (**Fig. 4**). Human-enriched set also include other six genes encoding virulence factors, namely SsrA, a major transcriptional regulator of SPI-2, which is involved in intracellular virulence (+3.2%);^30^ the SPI-4 encoded SiiA and SiiF, an accessory protein and a component of the SiiCDF type I secretion system, respectively, involved in adhesion and intestinal colonization (+2.5% and +20.6%);^31^ SopD2, an effector protein secreted by the SPI-2 type III secretion system and involved in intracellular survival and manipulation of host-cell trafficking (+5.6%);^32^ RtsA, a transcriptional activator of SPI-1, which is involved in epithelial cell invasion (+11%);^33^ Ssek1, involved in survival inside macrophages.^34,35^

**Figure 4.**
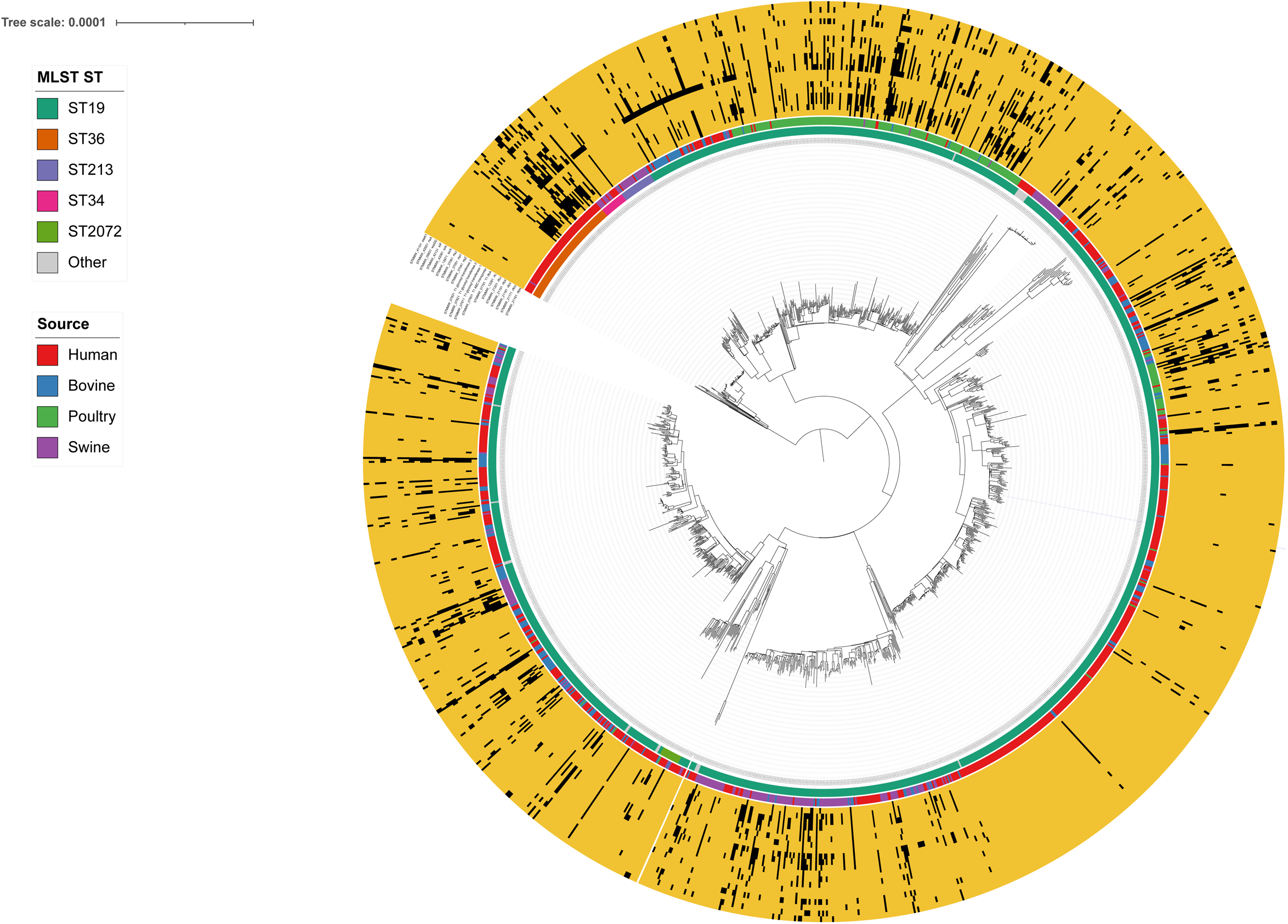
Distribution of the human-enriched genes across the *S.* Typhimurium phylogeny. Recombination-filtered maximum-likelihood core-SNP phylogeny of the 1,398 genomes, rooted on ST36. The inner ring indicates the MLST sequence type and the outer ring the isolation source. The heatmap shows the functional status of the human-enriched genes — the *rfb* O-antigen genes, the T1-antigen region, the *rfa* lipopolysaccharide core genes and the virulence genes *ssrA*, *siiA*, *siiF*, *sopD2*, *rtsA* and *sseK1* — coloured where a functional copy is present and black where the gene is absent or truncated

Genomes lacking the described human-enriched genes in functional form do not cluster into specific evolutionary lineages, rather, they appear homogeneously widespread across the ST19 population (**Fig 4**). These results indicate that their loss/deleterious mutation does not become fixed in the population.

The human-depleted set was characterized by genes involved in carbohydrate utilization. Specifically, seven genes encoding the PTS glucose transporter subunit IIABC (STM2758) and the galactitol utilization gene cluster (STM3255- STM3260) were found.^36^ The frequency of these genes was, on average, 30% ± 0.10% lower in human- than in livestock-associated genomes. Moreover, the human-depleted set includes five virulence genes, namely *ratB* and *shdA*, located within the CS54 genomic island of S. Typhimurium associated with intestinal colonization and persistence (-32 ± 1%);^37^ *lpfD* and *stfF*, encoding components of fimbriae implicated in bacterial adhesion and colonization of host tissues (-31.3% and -3.2%);^38,39^ *pipB2*, encoding a type III secretion system effector protein involved in intracellular interactions with host cells and in the modulation of host cellular processes (-9.8%).^40^ Genomes lacking genes involved in carbohydrate utilization, as well as *ratB* and *lpfD*, tended to cluster within specific evolutionary lineages, suggesting that these genes were lost or inactivated early in the evolution of these lineages and subsequently maintained as lineage-specific traits (**Fig. 5**).

**Figure 5.**
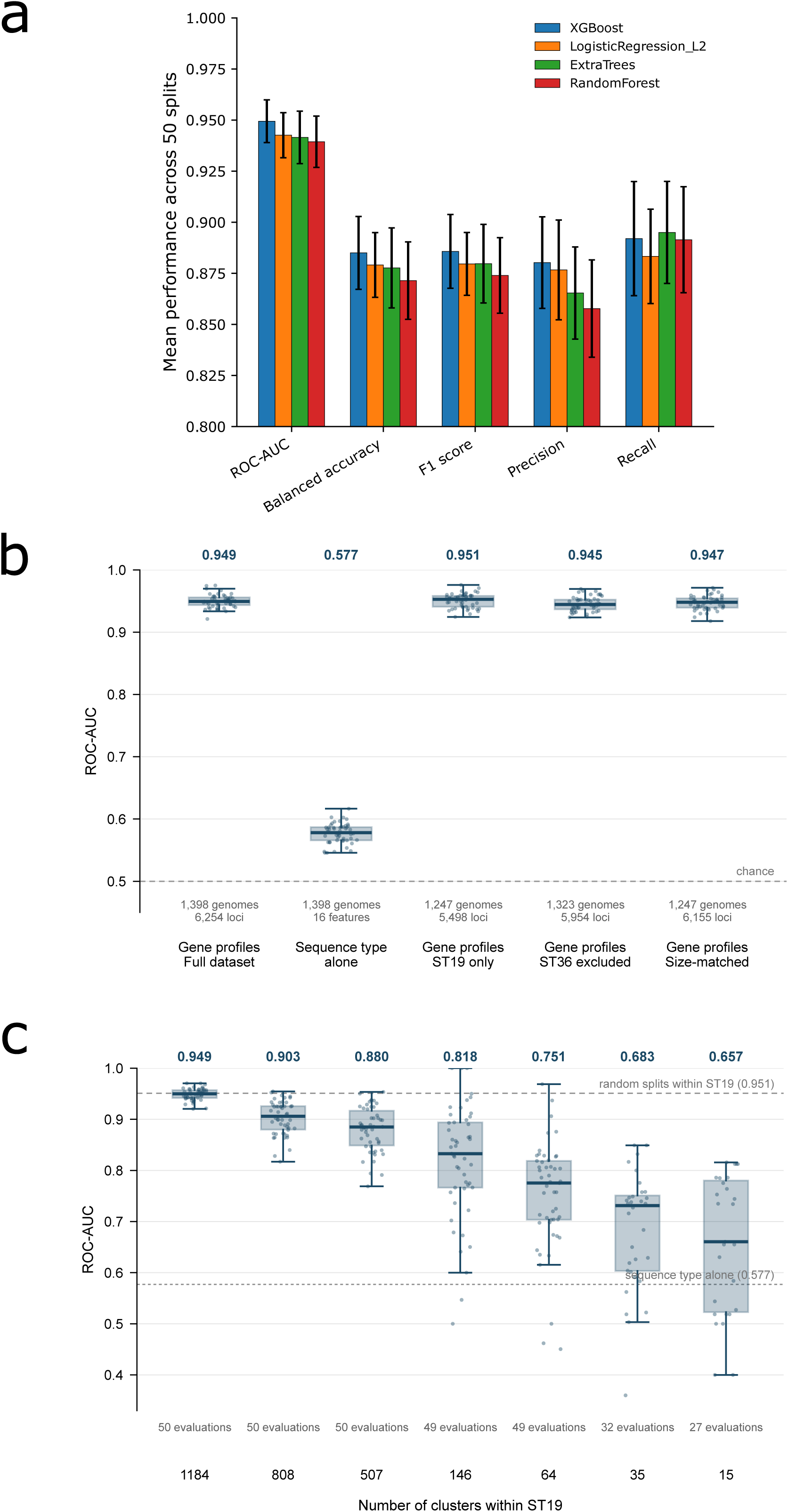
Distribution of the human-depleted genes across the *S.* Typhimurium phylogeny. Recombination-filtered maximum-likelihood core-SNP phylogeny of the 1,398 genomes, rooted on ST36. The inner ring indicates the MLST sequence type and the outer ring the isolation source. The heatmap shows the functional status of the human-depleted genes — the glucose and galactitol phosphotransferase-system transporters, the CS54 island genes *shdA* and *ratB*, and *lpfD*, *stfF* and *pipB2* — coloured where a functional copy is present and black where the gene is absent or truncated.

To investigate whether human-enriched and human-depleted genes tend to co-localise within the genome, their reference sequences were mapped to the *S.* Typhimurium LT2 genome and genes separated by no more than 10 kb were considered as belonging to the same cluster. Forty-five of the 97 human-enriched genes (46.4%) and all 36 human-depleted loci were successfully mapped to the reference genome (**Fig. 6a**). The human-enriched genes formed nine clusters containing 28 of the 45 mapped genes (62.2%), whereas the human-depleted genes formed three clusters containing 15 of the 36 mapped genes (41.7%); the two largest are shown in **Fig. 6c**. Compared with 20,000 random redistributions of the same number of genes along the chromosome, significantly more human-enriched genes were located within clusters than expected by chance (28 observed vs. 7.5 expected, *p* < 0.0001; **Fig. S4a**). Similarly, human-depleted genes were significantly more clustered than expected (15 observed vs. 4.9 expected, *p* = 0.0010; **Fig. S4b**).

**Figure 6.**
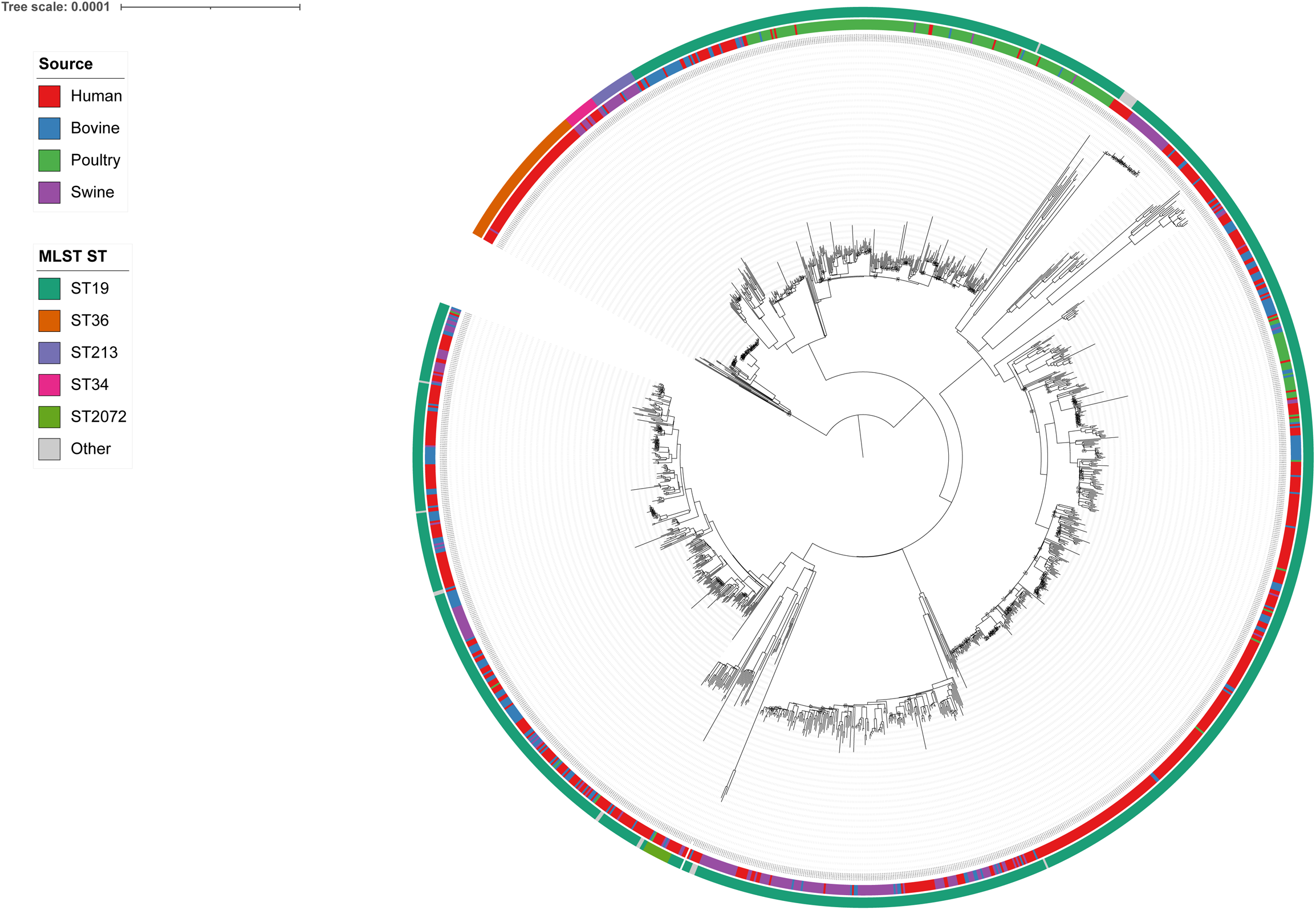
Genomic localisation of the ST19 human-enriched and human-depleted genes. (a) Positions on the LT2 chromosome (NC_003197.2) of the human-enriched and human- depleted genes. Tick blocks represent individual genes; shaded blocks represent clusters of two or more genes, labeled C1 to C9 (human-enriched) and C1 to C3 (human-depleted) in order of genomic position. **(b)** Gene maps of three human-enriched clusters and **(c)** of two human- depleted clusters. Arrows represent coding sequences oriented according to their strand. Filled arrows indicate human-enriched and human-depleted genes, whereas outlined arrows represent the other genes annotated in the region.

Regarding the human-enriched set, three of the four largest clusters included gene involved in LPS biosynthesis, specifically five genes of the *rfb* locus encoding the rhamnosyltransferase RfbN, the glycosyl transferase RfbU, the abequosyltransferase RfbV, the O-antigen transporter RfbX and the CDP-abequose synthase RfbJ; five genes of the T1-antigen locus encoding the O- antigen export system permease RfbA, a polysaccharide export ABC transporter and three glycosyl transferases; and three genes of the *rfa* (*waa*) locus encoding the 3-deoxy-D-manno- oct-2-ulosonate III transferase RfaZ, the LPS core heptose(II) kinase RfaY and the glucosyltransferase RfaJ (**Fig. 6b)**.

Among the human-depleted clusters, one corresponded to the galactitol utilisation operon, encoding the galactitol phosphotransferase-system (PTS) subunits IIA, IIB and IIC (STM3255, STM3258- STM3260), a fructose PTS subunit IIA (STM3256) and a tagatose-bisphosphate aldolase (STM3257). Another human-depleted cluster comprised the virulence genes *shdA* and *ratB*, both encoded in the CS54 intestinal colonisation island (**Fig. 6c)**.

## Discussion

An important question arising from the epidemiology of human salmonellosis is whether the strains causing human disease represent a random subset of the *Salmonella* population circulating in livestock and transmitted through contaminated livestock-derived foods, or whether they reflect a non-random subset enriched for specific genetic features that facilitate human disease. To address this question, we focused on *S*. Typhimurium, a major cause of human salmonellosis, and assembled a balanced dataset of approximately 1,400 genomes from human and livestock-associated isolates for comparative analysis.

The dataset includes only genomes from isolates collected within the same geographic region (North America), to compare strains associated with human disease with those circulating in livestock in the same area, which may serve as a proxy for the *S.* Typhimurium population to which humans are likely to be exposed.

The dataset was first analysed through core SNP phylogeny that did not clearly separate human-associated from livestock-associated genomes. Therefore, we evaluated whether supervised machine-learning approaches could discriminate between the two groups based on their functional gene presence/absence profiles. The results of four supervised machine- learning algorithms indicated that human-associated *S.* Typhimurium genomes were distinguished from livestock-associated genomes with consistently high predictive accuracy based on the presence or absence of functional genes. This supports the hypothesis that human-associated strains represent a non-random subset of the *Salmonella* population circulating in livestock with distinct genetic features.

This discrimination was not substantially influenced by the phylogenetic structure of *S.* Typhimurium at the ST level, but part of the predictive signal was associated with population structure within the most common ST19, where distinct lineages showed predominant associations with specific hosts and were already characterized for their distinct genomic content.^6^

Nevertheless, among the 1,774 genes contributing to the prediction, we identified genes whose frequencies differed significantly between ST19 human-associated genomes and each of the three animal hosts, indicating a consistent association with human disease.

Notably, among the genes more frequently present in ST19 human-associated genomes, we identified several genes involved in the biosynthesis of the LPS core, O-antigen and T1 antigen. Remarkably, two of these genes (*rfbV* and *rfbX*) were also enriched in human-associated genomes belonging to ST2072, suggesting that their association with human disease extends beyond the ST19 lineage. These findings strongly suggest that maintaining an intact LPS biosynthetic machinery may contribute to the ability of *S*. Typhimurium to cause disease in humans, consistent with the central role of LPS in multiple steps of host-pathogen interaction, including intestinal colonization, serum killing resistance, induction of inflammatory response, invasion, intracellular replication and resistance to killing by macrophages. A similar consideration may apply to the T1 antigen, although its precise biological function remains to be fully elucidated.^29,41–44^ The consistent enrichment of five of its eight biosynthetic genes in human-associated genomes across all three animal comparisons suggests a potential role for T1 in human salmonellosis.

The human-enriched genes also included six genes encoding known virulence factors (SsrA, SiiA, SiiF, SopD2, RtsA, and SseK1), supporting their potential role in human pathogenicity.

The higher frequency of virulence genes in human- than in livestock-associated genomes likely reflects the nature of the isolates from the two sources. Human-associated *Salmonella* isolates are predominantly obtained from microbiologically confirmed cases of salmonellosis, and therefore largely represent strains associated with human disease. In contrast, livestock- associated isolates may be recovered not only from clinically affected animals but also from apparently healthy or subclinically infected animals through routine surveillance and monitoring programs. Thus, the animal collection may encompass a broader spectrum of *S.* Typhimurium isolates, including those capable of colonizing or persisting in livestock without causing overt disease. In these isolates, successful colonization and persistence may not require the full repertoire of virulence factors associated with disease, potentially allowing the loss or inactivation of genes whose functions are more relevant to pathogenicity than to asymptomatic host persistence.

Interestingly, other virulence genes (*ratB*, *shdA, lpfD*, *stfF*, and *pipB2*) were instead less frequently present in functional form in human- than in livestock associated genomes. This finding suggests that human-associated genomes are not characterized by a general enrichment of virulence-associated genes; rather they appear to rely on specific genetic determinants. One possible explanation is that the classification of these genes as virulence factors does not necessarily imply a direct role in human infection, since their functions may have been characterized in particular host species or experimental models. Given the variation in host-pathogen interactions across different species, the contribution of these genes to virulence could be host-dependent.

Among the genes less frequently present in human- compared to livestock-associated genomes, we also identified those involved in the galactitol utilization pathway. As *in vivo* experiments demonstrated that the ability to metabolize galactitol contributes to *S.* Typhimurium intestinal colonization of food-producing animals, the higher frequency of these genes in livestock-associated genomes suggests that galactitol utilization may confer an advantage for persistence in livestock but may not be required for human disease.^36,45^ Consistent with this hypothesis, we previously found genes involved in the galactitol utilization pathway to be enriched in attenuated *Salmonella* serovars, which are frequently isolated from livestock but contribute only marginally to human disease.^46^ The recurrence of this genomic pattern across phylogenetically distinct *Salmonella* lineages suggests that it may reflect a form of convergent evolution associated with adaptation to livestock rather than to the human host.

However, the loss or inactivation of genes depleted in human-associated genomes was primarily associated with a specific ST19 sublineage that was predominantly composed of human-derived isolates. Therefore, it remains unclear whether this genetic signal merely reflects the evolutionary specificity of this lineage or whether these genes are genuinely dispensable for causing disease in humans. This result further supports the notion that part of the predictive signal was driven by genetic features characteristic of specific lineages that were predominantly associated with particular host sources, highlighting the importance of interpreting machine- learning results considering the underlying phylogenetic structure of the dataset.

Overall, our findings demonstrate the potential of machine-learning approaches to identify genetic features associated with human disease from naturally occurring variation in *S.* Typhimurium populations. Notably, among genes consistently enriched across all three human– animal comparisons, we identified not only known virulence genes but also several genes with currently unknown functions. Their reproducible association with human-associated genomes supports their consideration as promising candidates for further investigation and raises the possibility that previously uncharacterized genetic determinants may contribute to human pathogenicity.

## Author contributions

Conceptualisation: GM, SP and MT. Data curation: GM and MT. Formal analysis: GM. Funding acquisition: ES and MT. Investigation: GM and MT. Methodology: GM, ES, and MT. Project administration: ES and MT. Resources: ES. Software: GM. Supervision: ES, SP and MT. Validation: GM, SP and MT. Visualisation: GM. Writing original draft: GM, ES, SP and MT. Writing, reviewing, and editing: all authors.

