## Supplementary figures for "Genomic signatures distinguish isolates of *Salmonella* Typhimurium from humans and livestock"

Giovanni Mericia,b*, Erika Scaltritia, Stefano Pongolinia, Martina Tambassia,c

^a^Center for Genomic Epidemiology – Research and Education Department, Istituto Zooprofilattico Sperimentale della Lombardia e dell'Emilia-Romagna, 43126 Parma, Italy

^b^Department of Chemistry, Life Sciences and Environmental Sustainability, University of Parma, Parco Area delle Scienze 11/a, 43124 Parma, Italy

^c^Department of Surgical, Medical, Dental and Morphological Sciences with Interest in Transplant, Oncological and Regenerative Medicine, University of Modena and Reggio Emilia, 41124 Modena, Italy

This file contains:

Supplementary Fig. S1

Supplementary Fig. S2

Supplementary Fig. S3

Supplementary Fig. S4

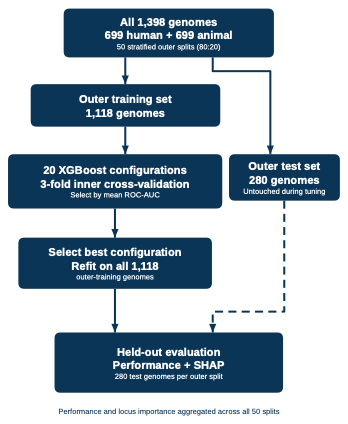

**Supplementary Figure S1. XGBoost and SHAP workflow.** XGBoost was tuned by three-fold cross-validation within 50 stratified train/test splits, with performance and SHAP values calculated on held-out genomes.

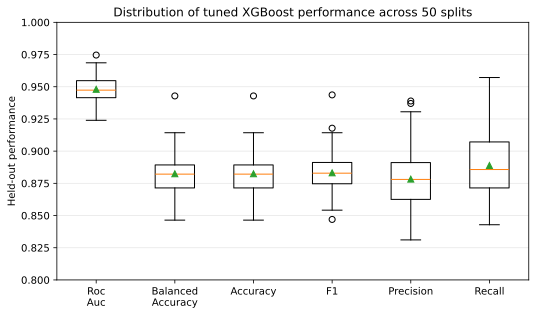

**Supplementary Figure S2. Held-out performance of the tuned XGBoost models across 50 stratified outer splits.** Each box summarises 50 values, one per outer split (280 held-out genomes each). Boxes span the interquartile range, orange lines the median, green triangles the mean.

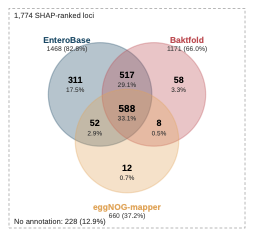

**Supplementary Figure S3. Overlap of the functional annotations obtained for the 1,774 SHAP-ranked loci by the three approaches.** Loci were counted as annotated when the product was other than "hypothetical protein" (EnteroBase, Baktfold) or a COG category other than S was assigned (eggNOG-mapper); loci outside the circles received no annotation from any approach.

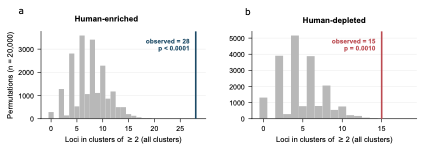

**Supplementary Figure S4. Permutation test of the spatial clustering of the ST19 discriminant genes.** Null distributions obtained from 20,000 permutations in which the same number of human-enriched genes **(a)** and human-depleted genes **(b)** were uniformly distributed at random along the LT2 chromosome and subsequently grouped using the same 10-kb criterion. The histogram shows the expected number of genes falling into two or more clusters, summed across all clusters, and the vertical line marks the observed value.
